# Melanopsin ganglion cells in the mouse retina independently evoke pupillary light reflex

**DOI:** 10.1101/2024.05.14.594181

**Authors:** Jeremy Matthew Bohl, Abdul Rhman Hassan, Zachary J. Sharpe, Megi Kola, Mahnoor Ayub, Yamini Pandey, Angela Shehu, Tomomi Ichinose

## Abstract

**Purpose:** The pupillary light reflex (PLR) is crucial for protecting the retina from bright light. The intrinsic photosensitive ganglion cells (ipRGCs) in the retina mediate the PLR, which directly sense light and receive inputs from rod/cone photoreceptors. Previous work used genetic knockout mice to reveal that rod/cone photoreceptors drive transient constriction, and ipRGCs drive the sustained component. We acutely ablated photoreceptors by a chemical injection to examine the role of rod and cone photoreceptors in PLR.

**Methods:** PLR and the multiple electrode array (MEA) recording were conducted with C57BL6/J (wildtype: WT) and Cnga3^-/-^; Gnat1^-/-^ (rod/cone dysfunctional) mice. n-Nitroso-n-methylurea (MNU) was applied to C57 mice by intraperitoneal injection, and PLR was conducted after 5-7 days of injection. Three different light levels (mesopic, low photopic, and high photopic) were tested. Immunohistochemistry was conducted using the anti-Gnat1 and anti-melanopsin antibodies with DAPI.

**Results:** PLR was induced by all light levels we tested, and the level of constriction increased as the light level increased. After the MNU injection, PLR was not induced at mesopic light stimulus, but was fully induced by high light. The level of PLR was identical between WT and MNU mice, suggesting that ipRGCs fully contributed to the PLR at this light level. Immunohistochemistry revealed that photoreceptors were ablated by the MNU injection, but ipRGCs were preserved. The MEA recording revealed that a population of ipRGCs generated fast and robust spikes in MNU-injected retinal tissues in *ex vivo*.

**Conclusions:** Contrary to previous observations, our results demonstrate that ipRGCs are the major contributor to the PLR induced by high light.

## Introduction

The pupillary light reflex (PLR) is induced by illumination to the eyes, constricting the pupil and protecting the eyes from excess light exposure. The PLR pathway starts from the retina, going through the optic nerve, the olivary pretectal (OPN) and Edinger-Westphal nuclei in the midbrain, and returning to the ciliary ganglion in the orbit, constricting the pupil. Because the PLR is easily examined, it has been broadly used in the clinic for diagnosing eye and brain diseases, such as glaucoma and brain injury. (La Morgia et al., 2018; Pinheiro and da Costa, 2021)

A specific type of retinal neuron conveys the PLR. Lucas et al. (2001; 2003) reported that the rod/cone degeneration mice, *rd/rd*, retain a robust PLR at high irradiances. However, when the melanopsin gene was also eliminated, PLR was absent (Lucas et al., 2003). These results indicate that the melanopsin-expressing cells directly sense light and induce PLR. These cells are categorized as intrinsically photosensitive retinal ganglion cells (ipRGCs), crucial for non-image-forming vision, including PLR and circadian photoentrainment (Hattar et al., 2003; Guler et al., 2008; Aranda and Schmidt, 2021). The rd/rd mice do not show the PLR in scotopic to mesopic light (Lucas et al., 2001; Lucas et al., 2003), suggesting that rod/cone photoreceptor inputs to ipRGCs induce PLR at the lower light conditions, and melanopsin is responsible for the high irradiances PLR.

Recent reports used photoreceptor mutant mouse models to dissect the roles of rod/cone photoreceptors and ipRGCs in PLR. In rod/cone dysfunctional mouse and human models, the PLR was evoked at high irradiances but with significantly slow onset compared to the control subjects (Gooley et al., 2012; Kostic et al., 2016). The onset delay was also observed in mGluR6-KO mice, where rod/cone signaling is not mediated onto ON bipolar cells and downstream ganglion cells (Beier et al., 2022). On the contrary, in the opn4^-/-^ mice, in which melanopsin is dysfunctional, PLR was induced across a wide range of irradiances; however, the pupil constriction occurred only transiently during the light exposure (Zhu et al., 2007). These reports indicate that rod/cone photoreceptors are crucial for the PLR at all light levels to induce rapid onset of the PLR, and ipRGCs shape the sustained PLR phase.

Most of the PLR studies have been conducted using transgenic mice, and we wondered whether those mice might have compensatory effects or degenerative changes. We used N-methyl-N-nitrosourea (MNU) to ablate rod/cone photoreceptors in a short span of 5-7 days (Smith and Yielding, 1986; Smith et al., 1988), a method that minimizes the potential for compensatory effects or degenerative changes. This approach allowed us to examine the roles of rod/cone photoreceptors and ipRGCs in the PLR, revealing unexpected features of ipRGCs.

## Methods

### Mouse Information

All experimental procedures with animals were approved by the Institutional Animal Care and Use Committee at Wayne State University (IACUC 20-10-2909 & 23-11-6310). Experiments were performed in accordance with the ARVO Statement for the Use of Animals in Ophthalmic and Visual Research. Wildtype (WT) (C57BL/6J; #000664, Jackson Laboratory, ME, USA) and rod/cone double knockout (Cgna3-/-, Gnat1-/-) mice (dKO, gift from Dr. Samar Hattar) ranging from 1-6 months in age of both sexes were used in this study. Using some WT mice, the n-nitroso-n-methylurea solution (MNU, 62.5 ng/g, HY-34758, MedChem Express, NJ, USA) or saline was injected intraperitoneally (i/p.), and the pupillometry or MEA recordings were conducted five to seven days after the injection.

### Pupillometry

C57 mice were dark adapted for over an hour on the day of the behavior recordings prior to any experiments and all PLR experiments were performed between 3-6 Zeitgeber time. For all recordings, mice were unanesthetized and mechanically restrained. To avoid a stress response, mice were handled in the experiment room prior to PLR testing. Mice were acclimated overnight in the experimental room before recording days.

For each PLR measurement, a mouse eye was observed by a stereo microscope equipped with a fixed-focus video camera (Teledyne Lumenara 4000) under illumination from an 850 nm infrared LED (ThorLabs). 500 nm (green) LEDs were used to induce the PLR in dark adapted mice. Light levels were measured and confirmed using a photometer (International Light Technologies, Peabody MA). PLR recordings were carried out across three light levels: Low (mesopic 5.34x10^4^ photons/μm^2^/s), Medium (photopic 2.95x10^5^ photons/μm^2^/s), and High (photopic 3.72x10^6^ photons/μm^2^/s).

During PLR recording, a five second baseline period was captured to measure the pupil before light onset. Subsequently the recording eye was flashed with green LEDs (500 nm) for ten seconds to measure transient pupil constriction. Recording continued until the pupil diameter stabilized after light offset. Pupils were recorded for at least 20s following light onset. Video recording of the PLR were captured at ∼25 frames per second (FPS) (NorPix 9;NorPix Quebec, CA).

### Retinal preparation

The experimental techniques were similar to those described previously. (Ichinose et al., 2014; Ichinose and Hellmer, 2016). Briefly, mice (28–60 days of age) were euthanized using carbon dioxide and cervical dislocation. Using a stereomicroscope equipped with infrared eyepieces, the retina was isolated and wholemount preparations were made under dark conditions. For some experiments, the wholemount preparation was placed onto a piece of filter membrane (HABG01300, Millipore) and cut into slice preparations (250 µm thick) using a hand-made chopper. Retinal dissections were performed in HEPES-buffered solution containing the following (in mM): 115 NaCl, 2.5 KCl, 2.5 CaCl_2_, 1.0 MgCl_2_, 10 HEPES, 28 glucose, adjusted to pH 7.37 by NaOH. The dissection medium was continuously oxygenated. The retinal preparations were stored in an oxygenated dark box at room temperature.

### Immunohistochemistry

We used our standard procedure for immunohistochemistry (Farshi et al., 2016). After fixing the retinal wholemount preparations using 4% paraformaldehyde for 60 minutes, sections were blocked in a solution containing 10% normal donkey serum (NDS) and 0.5% Triton-X in PBS (PBS-T) for 1 hour at room temperature (RT). Melanopsin antibody (AB-N39, Advanced Targeting Systems, CA, USA) was diluted (1:5000) in 3% NDS and PBS-T and incubated with the tissue for 3 days in 4 °C. Then, tissues were incubated with Alexa568 donkey-anti-rabbit (Invitrogen; A10042) for 2 hours. Gnat1 antibody (PA5-28336, Thermo Fisher) was diluted (1:1000) and incubated for overnight at RT. DAPI (1:10000, Sigma, Co.) was added and incubated for 20 minutes. Stained retinas were mounted on the slide glass and viewed with a confocal microscope (TCS SP8, Leica). The z-step for stack images was 0.3 μm.

Melanopsin and DAPI cell counting was performed using confocal microscope images. Z-stacks of the ganglion cell layer were max projected using ImageJ (NIH) to improve image quality and reduce background noise. For each condition, four quadrants measuring at least 140 x 140 μm were manually annotated from each image. Cell counts were averaged from all regions of both retinas (n= one mouse) for statistical analysis. A one-way ANOVA using Tukey’s Method post-hoc comparisons were used to compare DAPI and melanopsin cell counts between saline, MNU injected, and dKO mice. A p-value < 0.05 was considered significant.

### Multi-electrode array (MEA)

WT, MNU, and dKO mice were dark adapted overnight and retinal wholemount preparations were prepared under infrared light using infrared viewers. The retinal preparation was placed with the ganglion cell side down onto the Accura HD-MEA chip (4,096 recording electrodes arranged in a 3.8 mm x 3.8 mm area multielectrode array) for the BioCAM DupleX system (3Brain, Switzerland). The tissue was anchored down by placing a flat filter paper on the photoreceptor side and the filter paper was held down by a square platinum anchor. The preparation was continuously perfused with oxygenated AMES’ medium at a rate of 3–7 ml/min and maintained at 30-34°C with a temperature controller (Multi Channel Systems, ALA Scientific). In some recordings, a cocktail of glutamate receptor antagonists was applied to block photoreceptor synaptic inputs to third-order neurons: an mGluR6 antagonist, AP4, an ionotropic glutamate antagonist, CNQX, a kainite antagonist, ACET, a NMDA antagonist, AP5.

The raw data of the MEA recording were analyzed using the Brainwave 5 software (3Brain, Switzerland). A preset spike detection configuration optimized for retinal wholemounts was utilized to detect spiking activity. Afterwards, spikes were sorted into individual spiking units that constitute a single ganglion cell. This isolates any spiking cells sharing the same probe and also to exclude any signals not produced by the retinal tissue. Spike sorting and clustering involved the following features: the pre-peak waveform duration was set at 0.5 ms, while the post-peak waveform duration was set at 1.0 m_5_ s. Feature extraction utilized principal component analysis. Clustering was performed using Gaussian mixture models, with a minimum of two and a maximum of three spikes required for clustering. Outliers were discarded and duplicates were discarded if they shared more than 40% overlap in spike times. All retinal spike recordings were binned in 0.3 ms intervals and were normalized to the average dark-adapted baseline spiking rate in the first 10 s prior to light onset. To analyze the baseline firing rate units for each condition, units were grouped according to their spike rates during each interval, forming groups ranging from 0 to 30 spikes. The count of units within each spiking group was then divided by the sum of units in all other spiking groups, yielding percentages per unit plotted against the number of spikes. This process was carried out for each mouse type and subsequent comparisons were made.

ANOVA analysis was utilized to compare these datasets, with n= the number of retinas for each condition and a p-value <0.05 was considered significant for this analysis.

### PLR Video Analysis

Videos of the PLR were analyzed using AIVIA software (Leica) to track pupil diameter during the entirety of the video. The pupil diameter was measured in every video frame (∼25 frames/second) and used to generate the PLR response over time traces. Individual eye recordings were only excluded if the mouse moved out of camera view during the ten second light flash. All recordings were resampled to exactly 25 FPS after pupil analysis to allow for average traces of each condition to be generated. For each recording, all measurements were normalized to the average dark-adapted pupil size across the 5 s immediately prior to light onset. Dark-adapted pupil diameters did not differ between mutant and control mice (one-way ANOVA post hoc, p > 0.5 between mutant mouse lines and controls). No differences were detected during ANOVA analysis between pre-MNU, pre-saline, and post-saline injection peak PLR constriction (data not shown p>0.05). As such, pre- and post- MNU injection curves were compared for analysis. For the analysis of the transient constriction during high light conditions, raw traces were fitted with a single component exponential function (R^2^ > .90, SigmaPlot14.5, Systat). A paired Student’s *t-test* was used to compare pre- and post- MNU injection tau values from every fitted curve. The peak constriction of pre- and post- MNU responses were manually extracted from the raw response over time curves and compared using a paired Student’s *t-test*. For all statistical measures n= the number of recorded eyes or number of mice and p-value < 0.05 was considered significant.

## Results

### Pupil light reflex from C57 wildtype, MNU, and rod-cone knockout mice

We conducted PLR from mouse eyes to examine retinal cellular contributions. The PLR was evoked in C57 wildtype (WT) mouse eyes by green light stimuli of three different levels. Mice were dark-adapted and all procedures were conducted in dim red-light conditions. In response to a low light flash for 10 seconds (10^4^ photons/µm^2^/s), pupils slightly constricted and recovered (Figure 1A, Low). As irradiance increased, pupil constriction was faster and more robust (Figure 1A, Medium and High, 10^5^ and 10^6^ photons/µm^2^/s, respectively). Multiple mouse eyes exhibited similar constriction patterns throughout PLR recordings (Figure 1B, WT, n=38 eyes from 19 mice).

**Figure 1.**
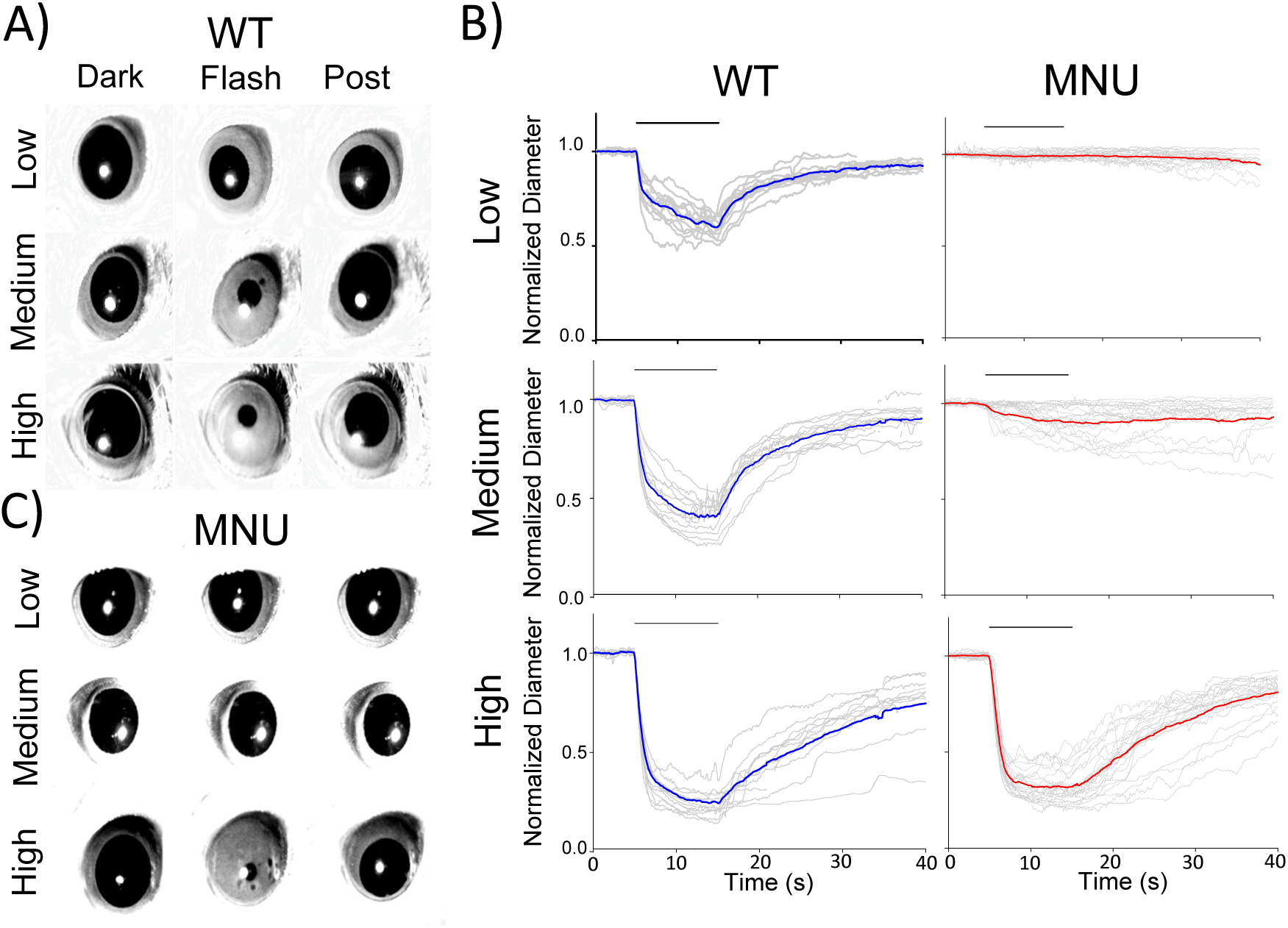
The pupillary light reflex (PLR) in WT and MNU-injected mice: A) Single frame images from PLR recordings for WT control mice. Pupil images before the light stimulus (Dark), during the light stimulus (Flash), and the recovery phase (Post) are presented for each light intensity. B) Normalized pupil diameters were plotted as a function of time. Individual eyes (grey traces) and average pupil constriction for WT (blue) and MNU (red) were overlaid. C) A set of representative pupil images for MNU-injected mice. PLR was barely detected in response to low and medium light stimuli. In contrast, high light-evoked PLR is similar to WT eyes.

The PLR is mediated by ipRGCs (Guler et al., 2008), which sense light directly through melanopsin (Lucas et al., 2003), as well as receiving photoreceptor synaptic inputs. To examine the differential roles of melanopsin and photoreceptors in the PLR, we conducted MNU *i.p.* injection into WT mice, which has been shown to eliminate photoreceptors in several days (Smith and Yielding, 1986; Smith et al., 1988; Wang et al., 2021).

Five to seven days after MNU injection, we measured the PLR (Figure 1B-C). The low-level stimulus did not evoke the PLR in MNU mice, suggesting that rods and cones are essential to PLR during mesopic light conditions. Medium light stimuli evoked a smaller constriction than WT eyes with slower onset and offset (Figure 1B-C, medium). These results are consistent with previous observations, showing ipRGC’s low sensitivity and sluggish melanopsin responses (Gooley et al., 2012; Kostic et al., 2016; Beier et al., 2022). We then increased the stimulus light level to high irradiance. Unexpectedly, PLR was evoked fully and rapidly, identical to the WT PLR (Figure 1B, High).

We examined the high light-evoked PLR onset speed and latency in WT and MNU eyes (Figure 2). The MNU eyes showed a longer latency from the stimulus to the PLR onset than those in WT eyes (Figure 2A-B, n=5 WT and n=5 MNU, p<0.05, unpaired Student’s *t-test*). However, the speed of PLR constriction was faster in MNU than in WT eyes (Figure 2C-D, p<0.05, paired *t-test,* n=12 mice). The level of constriction was the same between WT and MNU eyes (Figure 2E, p=0.94). These analyses confirmed that the high light-evoked PLR in MNU mice is equivalent to the WT eyes. Because the MNU eyes contain only ipRGCs, our results indicate that ipRGCs fully contribute to the PLR at this light stimulus.

**Figure 2.**
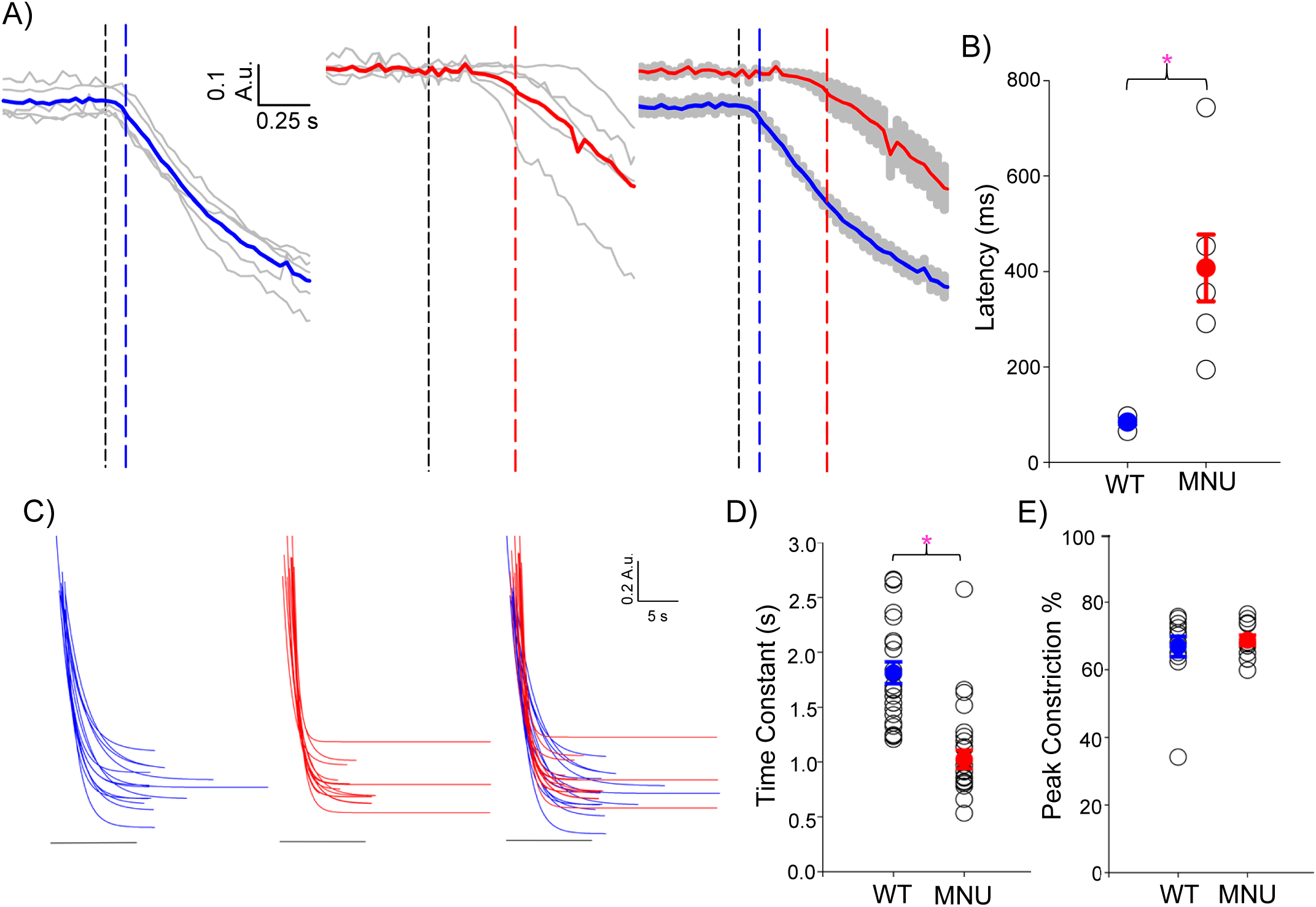
Temporal features of high light-evoked PLRs were equivalent between WT and MNU eyes: A) The initial phase of high light-evoked PLRs is displayed in a short time scale. The latency from the light stimulus onset (black dotted line) to the start of the PLR in WT eyes (blue dotted line) and MNU eyes (red dotted line) are shown. B) A summary graph shows the latency in WT and MNU eyes, which showed a significant delay (p<0.05, Student’s unpaired *t-test*). The means for WT and MNU are presented in blue and red. C) The onsets of high light-evoked PLRs were fit by single component exponential decay curves for WT eyes (blue) and MNU (red) injected mice (R2>0.90). D) A scatter plot showing each curve fit’s time constant (tau). MNU exhibited low tau, indicating a faster PLR constriction phase than WT PLR (p<0.05, n=12 mice). E) A scatter plot showing peak constriction showed no differences between WT and MNU mice (p= 0.07, n=12 mice).

Many previous reports used mutant mouse models to examine the role of ipRGCs in PLR, and show a become slower and smaller PLR in these mice (Gooley et al., 2012; Kostic et al., 2016; Beier et al., 2022)..To compare these with our MNU model, we used a mutant model in which rod-cone photoreceptors are dysfunctional (Cnga3^-/-^ & Gnat1^-/-^: dKO) and examined the PLR. As expected, low light minimally evoked the PLR, and medium light evoked small and slow PLR (Figure 3, Low & Medium). The high-light stimulus evoked a more robust response (Figure 3, High); however, the level of pupil constriction was smaller, and transient constriction speed was slower than PLRs in WT and MNU (p< 0.0001, Figure 3C). This observation with dKO mice was consistent with previous reports (Kostic et al., 2016; Beier et al., 2022). While both MNU and dKO mice only have the ability to use ipRGCs to respond to light, only MNU mice showed WT mouse-equivalent constriction in response to high light stimuli.

**Figure 3.**
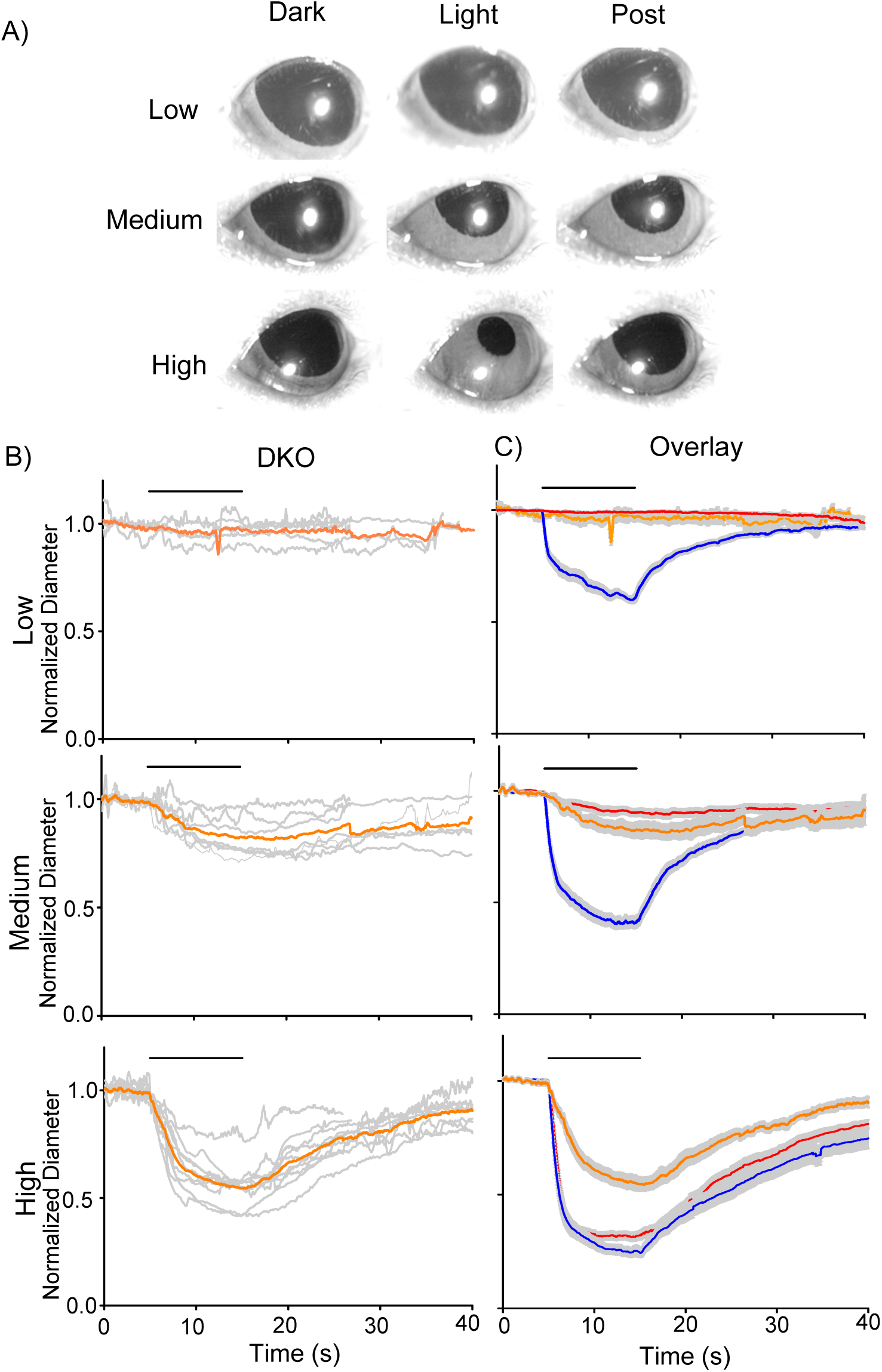
Double rod/cone knockout (dKO) mice showed an altered PLR compared to WT and MNU mice. A) A set of pupil images showing PLR from dKO mice. No PLR was detected by low light, and small constrictions were detected by medium to high light stimuli. B) Normalized pupil diameters over time were plotted from dKO mice (n=5). The individual dKO eyes were in grey and the average responses of all eyes were in orange. C) The mean PLRs for all three conditions are overlaid (WT: blue, MNU: red, and dKO: range) with the S.E.M. (grey).

### Photoreceptors were dysfunctional in MNU retinal tissue

The fast ipRGC response in the MNU tissues might be evoked by the remaining photoreceptors, which was shown previously (Jain et al., 2016). We conducted the tissue analysis to examine whether MNU injection removed photoreceptors. We first compared vertical retinal sections (Figure 4A). The MNU injection eliminated the outer retinal layers, from the outer plexiform layer (OPL) to photoreceptor outer segments (Figure 4A, n= 6 mice for saline, 12 mice for MNU). Retinal sections were taken from both central and peripheral retina, and show photoreceptor loss was uniform across the retina after the MNU injection.

**Figure 4.**
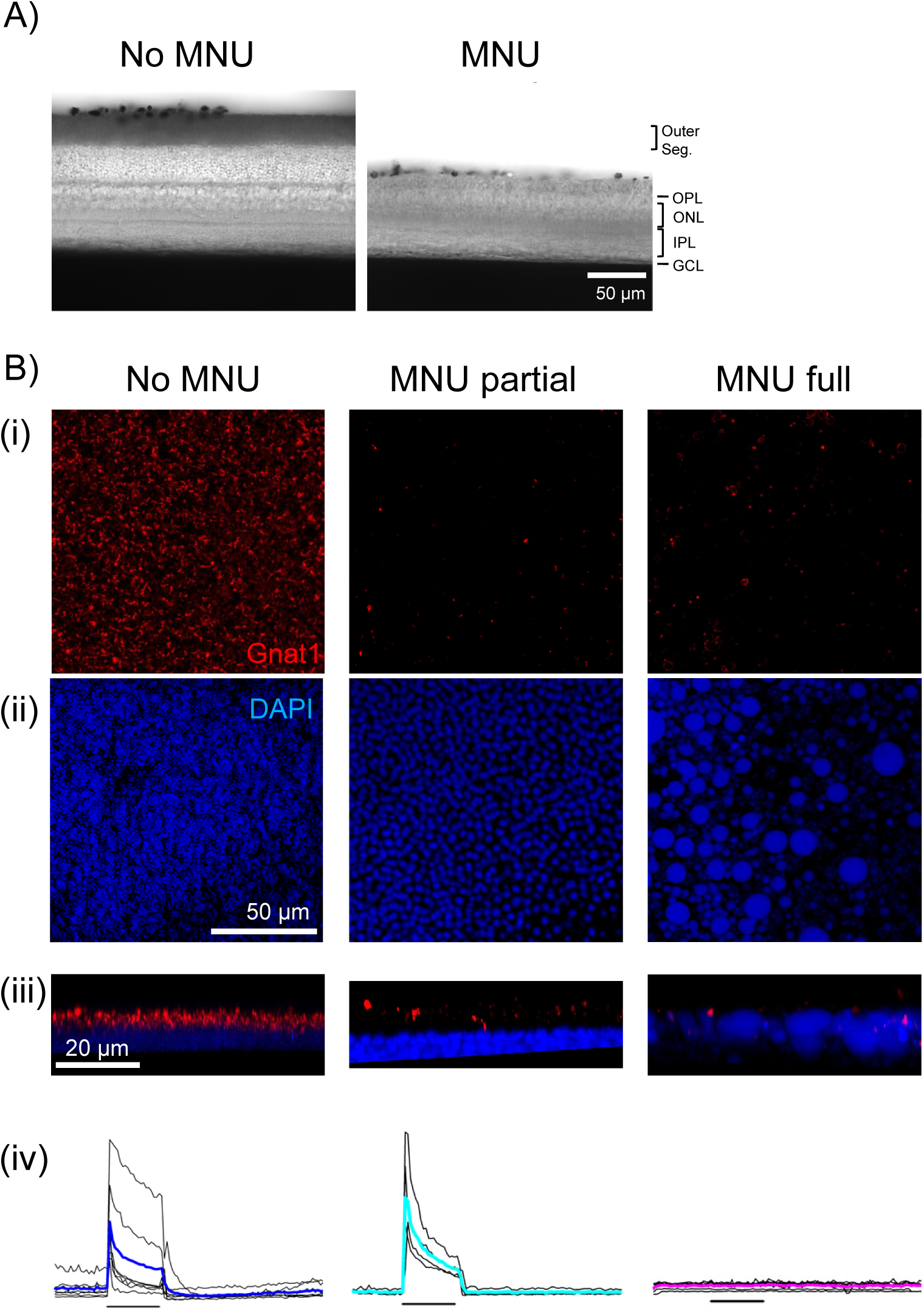
MNU selectively ablated rod/cone photoreceptors: A) DIC images showing retinal ve r t ic a l sections from WT (left) and MNU eyes (right). MNU-injected retinas showed a loss of the photoreceptor layer, including the outer segment and outer nuclear layer. The inner retinal layers remained intact. B) Immunohistochemistry images of the photoreceptor layer in wholemount tissues (i-iii). (*Left Column*) The WT mice without MNU revealed a massive Gnat1-labeled rod outer segment (red) and DAPI-labeled photoreceptor somas (blue). A lower panel (iii) shows a digitally rendered image of the side view. The lowest panel (iv) shows the MEA recording of ganglion cell spikes in response to a low light stimulus. (Middle *Column*) The MNU partial tissue revealed the decreased Gnat1 puncta with almost intact somas. The MEA recording showed low light-evoked light responses. (*Right Column*) In general, MNU injection almost abolished the Gnat1-labeled outer segment (i), which resided in the DAPI-labeled soma layer (iii). The somas were swelled and exhibited cobblestone-like irregular structures (ii). The side view revealed the swelled somas with the Gnat1 puncta intermingled. For these eyes, low light did not evoke ganglion cell spikes (iv).

However, degenerating photoreceptors could respond to light even after the majority of rods and cones are eliminated in rd10 mice (Ellis et al., 2023). We examined our MNU tissues using the anti-Gnat1 antibody and DAPI staining (Figure 4B i-iii). In WT tissues, Gnat1 stained rods’ outer segments and DAPI stained the photoreceptor somas (Figure 4B-i, “No MNU”). The side view of the layer by digital rendering, showing that somas and outer segments are polarized (Figure 4B-iii, “No MNU”). MEA recording revealed that many ganglion cells responded to a low light stimulus (Figure 4B-iv, “No MNU”).

When MNU was injected, the Gnat1 staining was significantly reduced (185,460 ± 31,900 rods/mm^2^, no MNU, n=3; 16,980 ± 46,300 rods/mm^2^, MNU partial, n=7, p<0.05, unpaired Student *t* test). However, in some cases, a low-light stimulus still evoked light responses similarly to control tissues in MEA recordings (Figure 4B-iv, “MNU partial”). For those cases, even though the Gnat1 staining was significantly reduced, their somas and the soma-outer segment polarization were preserved (Figure 4B-i-iii, “MNU partial”).

In general, MNU injection wiped out low light-evoked light responses in MEA recordings (Figure 4B-iv, “MNU full”) similar to low light responses recorded in dKO mice (data not shown), that are consistent with previous results. Gnat1 staining was significantly reduced (1,190 ± 700 rods/mm^2^, MNU full, n=4, p<0.05 vs. control, unpaired Student *t* test). Photoreceptor somas were swelled, and the soma-outer segment polarization disappeared (Figure 4B-ii, “MNU full”). Based on these observations, we considered MNU was fully effective when low light-evoked response disappeared. When light response or PLR was evoked by low light, we considered them MNU partial effect and excluded them from our MNU group.

MNU removed the photoreceptor layer, but did not affect the ganglion cell layer cells (GCL) (Figure 5A). We examined the GCL structure after the MNU injection by conducting immunohistochemistry with DAPI and the opn4 antibody staining, staining GCL nuclei and ipRGCs, respectively. We found that the number of entire GCL cells was similar between control and MNU tissues (Figure 5A-B, p=0.29, n=10 WT and n= 11 MNU). The number of ipRGCs varied among retinas, but the overall ipRGC number was the same between WT, MNU, and dKO tissues (Figure 5A & C, p=0.196, One-way ANOVA, n=12 WT, n=13 MNU, and n=10 dKO).

**Figure 5.**
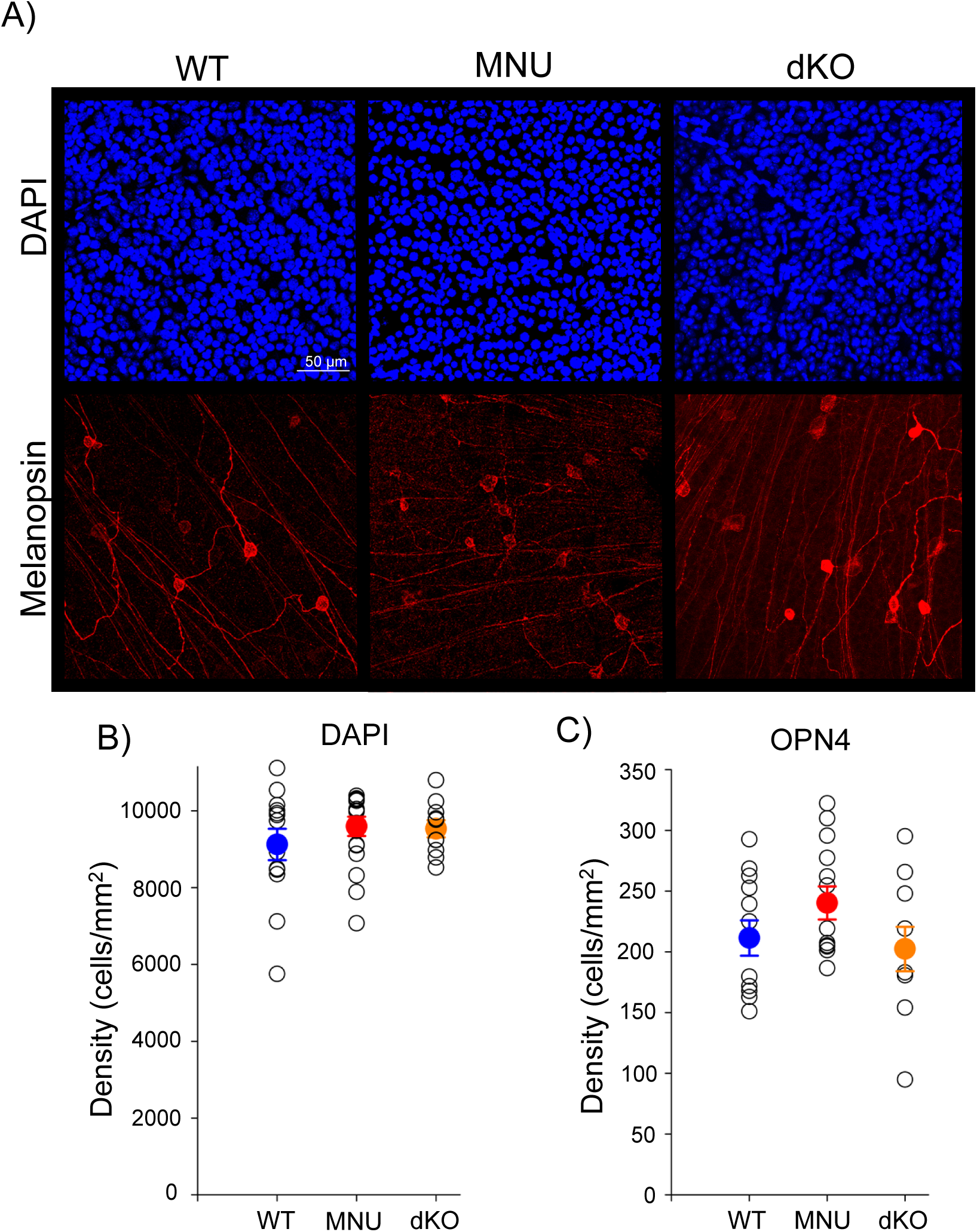
Ganglion cells were intact in MNU and dKO mice: A) Representative ganglion cell layer images after immunostaining with DAPI (blue) and OPN4 (ipRGCs, red) in WT, MNU, and dKO retinal tissues. B) The number of GCL somas by DAPI staining remained similar between WT, MNU, and dKO retinas (p=0.288, n=15 WT, n=17 MNU, and n=4 dKO mice, One-Way ANOVA with Tukey’s Method post-hoc comparisons). (C) The number of ipRGCs also remained constant in WT, MNU and dKO mice (p=0.196, n= 12 WT, n= 13 MNU, and n=10 dKO mice. One-Way ANOVA with Tukey’s Method post-hoc comparisons).

Furthermore, the number of GCL cells stained with DAPI remained consistent across experimental conditions (Figure 5, p=0.287, One-way ANOVA). These results suggest that a high light stimulus can induce a normal PLR in the absence of photoreceptor inputs to ipRGCs.

### ipRGC light-evoked response recording by MEA

To explore the physiological mechanisms underlying the differential PLR responses in WT, MNU, and dKO mice, we conducted MEA to record ipRGC light-evoked responses. We evoked ipRGCs using the high light stimulus, the same intensity used for PLR recordings. In WT tissues, most electrodes showed light-evoked responses immediately after light stimulus and/or light offset, indicating photoreceptor inputs. However, we identified spike units that lasted more than a second after the light shut-off which are thought to be ipRGCs (Figure 6A) (Qiu et al., 2005; Prigge et al., 2016). Consistent with melanopsin light transduction the spike responses of these units were not abolished by application of glutamate receptor antagonists (L-AP4, D-AP5, ACET and CNQX) to block photoreceptor synaptic transmission (data not shown).

**Figure 6.**
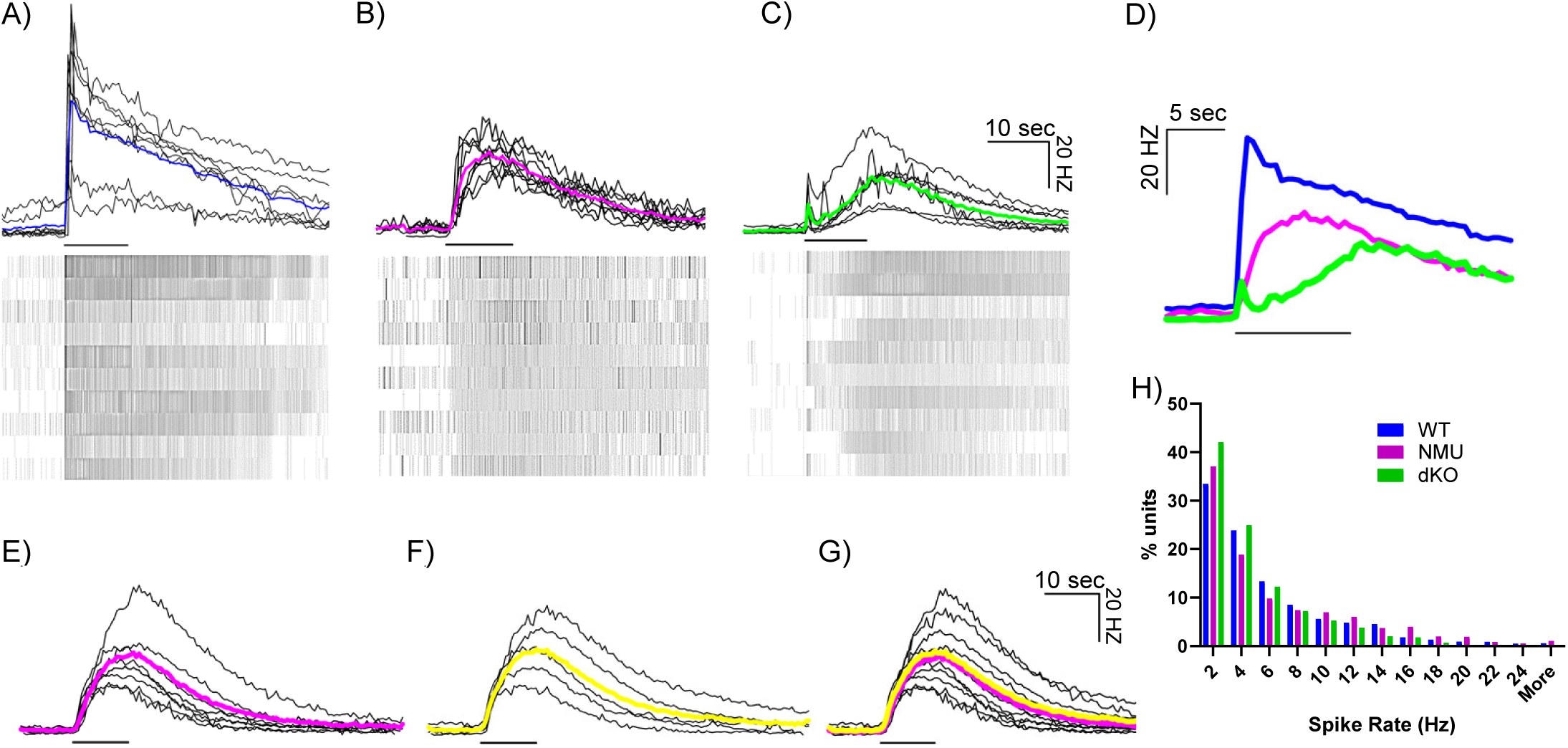
MEA recording from ipRGCs in three mouse types in response to high light stimuli: Individual trace in black indicates ipRGC units in WT (A), NMU (B) and dKO (C), and average traces are shown in color (WT: blue, n=10 tissues, MNU: magenta, n=9 tissues, dKO: green, n=5 tissues). Black lines under each trace indicates the 10 sec light stimulus timing. Below each trace there is a raster plot containing individual spikes of 10 representative ipRGCs from each respective tissue. (D) ipRGC average onset spike rate of the 3 different mouse types in response to high light stimuli. Three colors indicate three mouse strains. The average spike rate of MNU injected mice at high light intensity before (E) and During (F) perfusing Glut receptor blockers (L-AP4, D-AP5, ACET and CNQX) were perfused to block potential photoreceptor inputs. (G) An overlay of before and During perfusing Glut receptor blockers. (H) A histogram showing spontaneous spike rate from three different mouse types.

In MNU-injected retina tissues (n=10), photoreceptor-evoked, quick-onset light responses were not recorded. However, many spike units exhibited slow-onset long-lasting light responses, suggestive of ipRGCs. Low light stimulus did not evoke light responses in MNU retinas (Figure 4B iv). In contrast, when we stimulated MNU tissue with the high light, some units responded with fast spiking onsets (Figure 6B, 9.6% of spike units), potentially one of multiple ipRGC types (Aranda and Schmidt, 2021). While many other spike units were slower-onset (data not shown), these fast units exhibited comparable onset speeds and spike rates with WT spike units (Figure 6D). In order to further rule out any involvement of photoreceptors input in the fast response of MNU mice, we applied a cocktail of glutamate receptor antagonists to block synaptic transmission from photoreceptors to ipRGCs. We recorded spike rates before and after the perfusion of the receptor blockers (Figure 6E-F). When overlaid, no difference was detected (Figure 6G), indicating that photoreceptor-evoked inputs did not contribute to the response. This result along with the evidence of no response by a low light stimulus (Figure 4B) confirms that photoreceptor inputs were eliminated in MNU mice.

We recorded MEA with dKO mouse tissues to examine whether the population of fast-responding ipRGCs found in the MNU tissues at high light were present. In response to the light stimulus, an initial transient was evoked that quickly subsided followed by a slow onset long-lasting ipRGC responses were evoked (Figure 6C), suggesting reminiscent photoreceptor response (Allen et al., 2010). However, we did not find any fast onset ipRGC units that matched the kinetics of those in MNU retinas (Figure 6B). Figure 6D reveals that the dKO spiking response had a significantly slower ipRGC component onset compared to both WT and MNU tissues. These temporal results are consistent with our PLR results (Figure 1 & 3), suggesting that the different ipRGC light responses shape the PLR differences in these mice.

In some degenerative mouse models, spontaneous activities increase (Ye and Goo, 2007; Borowska et al., 2011), which may mask the light response of these cells and alter the threshold of light-evoked responses. We compared the baseline spontaneous spike rate in WT, MNU, and dKO retinal tissues. We measured the spike rate five seconds before any light stimuli and plotted the percentage of units in a histogram as a function of their spike rate (Figure 6H). Although the dKO tissue showed fewer units with high spike rates, there were no statistical differences among groups (n=5 tissues for each mouse strain, n= 190 – 542 units, p=0.999, one-way ANOVA). These results indicate that high spontaneous spike rates were not the reason ipRGCs did not show light responses in dKO mice, but ipRGCs might be remodeled or degenerative in these mice.

## Discussion

We examined the roles of rod/cone photoreceptors and melanopsin in PLR by MNU injection to eliminate photoreceptors within seven days. Although MNU mice showed reduced PLRs in response to low to medium light stimuli, a high light stimulus unexpectedly evoked fast and robust PLR, equivalent to WT mouse eyes. Then, ipRGC spikes were examined with the MEA in retinal tissues, which exhibited that approximately 10% of ipRGCs in the MNU tissue had fast onset. Our *in vivo* and *ex vivo* results indicate that the intrinsic light responses in ipRGCs are fully capable of inducing PLR at high-light stimuli, which contradicts the previous reports showing the pivotal role of photoreceptors in high-light-induced PLR.

### MNU injection and photoreceptors/ipRGCs

MNU is a DNA alkylating agent distributed widely in the environment as a potent carcinogen (Smith et al., 1988) and has been used to produce a retinal degeneration mouse model, which damages the photoreceptors without affecting inner retinal neurons (Kiuchi et al., 2002; Wang et al., 2021). Consistent with these previous reports, we observed that the retinal photoreceptor layers were eliminated only five days after MNU injection in all tested mice (Figure 4A). Although we observed that the photoreceptor outer segment was eliminated from the central to peripheral regions, reminiscent photoreceptors may still respond to a high-light stimulus. Jain et al. (2016) conducted immunostaining with s-opsin, m-opsin, and rhodopsin and observed reminiscent photoreceptors after 7 days of MNU injection. Furthermore, in rd10 mice, degenerating photoreceptors still respond to light even with a shortened outer segment (Ellis et al., 2023). Therefore, we examined reminiscent photoreceptors in the MNU tissue.

The MNU injection drastically reduced the photoreceptor layer and Gnat1-stained puncta (Figure 4). However, in some MNU tissues, photoreceptor-evoked responses were still evoked in response to low light stimuli, and photoreceptor somas were still observed (Figure 4B, “MNU partial”). In most MNU tissues, low light-evoked responses were not evoked (Figure 4B “MNU full”). In these tissues, we observed swelled somas near the OPL with small numbers of Gnat1 puncta hovering, indicating that photoreceptors were no longer functional in these retinas. We used the latter as MNU mice for both PLR and MEA recordings, ensuring that the fast and robust PLR and ipRGC responses in MNU mice were not attributable to photoreceptor responses.

Even though photoreceptors were eliminated in the MNU mice and dysfunctional in the dKO mice, ganglion cells were intact (Figure 5). Jain et al. (2016) observed the increased ipRGCs in the MNU tissue. The range of the ipRGC number was relatively large (Figure 5C), and the ipRGC number might be increased after the MNU injection in some cases. However, it is unlikely that increased ipRGCs in MNU eyes make new connections with postsynaptic targets in several days to boost the PLR. Taken together, we confirmed that MNU injection eliminated photoreceptors in several days without affecting ipRGCs.

### Contribution of rod/cone photoreceptors and ipRGCs to PLR

ipRGCs are crucial neurons for the non-image-forming vision, which also receive synaptic inputs from rod/cone photoreceptors, commonly the first-order neurons for the image-forming vision. The roles of these photosensitive cells in PLR have been investigated. Since rod photoreceptors are highly sensitive to light compared to cones and ipRGCs (Berson et al., 2002; Field and Rieke, 2002; Pang et al., 2004; Qiu et al., 2005), rods are crucial for low light-evoked PLR. A photoreceptor degeneration model, *rd/rd cl* mice, shows PLR only at high- but not low- to mid-irradiances (Lucas et al., 2001; Lucas et al., 2003). In these mice, PLR occurred at light levels equivalent to the operational range of the ipRGCs (Lucas et al., 2001). Our results with the MNU injection and dKO mice were consistent with those of the *rd/rd* mice with the threshold of 10^4^ photons/µm^2^/s (or 10^12^ photons/cm^2^/s), our low light stimulus (Figure 1-3).

The ipRGC’s unique property includes a high threshold to light stimuli and sluggish light responses (Berson et al., 2002; Dacey et al., 2005). Because of this slow nature, differential temporal roles of these photosensitive cells have been explored. Lucas et al. (2001) compared the high light-evoked PLR in wildtype and *rd/rd cl* mice and found that the onset of PLR was significantly delayed compared to wildtype. Kostic et al. (2016) used rod/cone dysfunctional mice (Cnga3^-/-^; Rho^-/-^ or Gnat1^-/-^) to reveal a small and slow PLR, which was consistent with the results of Beier et al. (2022), who used mGluR6-KO, Elfn1-KO, Cone-Cx36-KO mice in which photoreceptor transmission to third-order neurons are disconnected. A human study exhibited similar results, which investigated blind humans whose photoreceptors were dysfunctional but whose circadian rhythms were normal (Gooley et al., 2012). Complimentary, the opn4^-/-^ mice exhibit short-lasting PLR (Zhu et al., 2007). All these results demonstrate that photoreceptors are crucial for initiating the PLR even at the high light stimulus, and melanopsin serves for the sustained response during light exposure. Our dKO mice (Cnga3^-/-^; Rho^-/-^ or Gnat1^-/-^) show consistent results (Figure 3).

Nevertheless, our results with MNU mice did not show the same results. Because photoreceptors were eliminated in MNU mice, the onset time of high light-evoked PLR exhibited a significant delay compared to the WT mice (Figure 2A-B), consistent with the *rd/rd cl* mice (Lucas et al., 2001). However, in MNU mice, the constriction speed was faster than the WT mice (Figure 2C-D), generating overall equivalent PLRs to WT mice. Our MEA recordings support the PLR results. In MNU mice, we found that approximately 10% of ipRGCs generated spikes with fast onset, similar to the WT mice and significantly faster than those of dKO mice (Figure 6). This fast ipRGC light response is not likely attributable to the remodeling because the number of DAPI-stained ganglion cell layer cells and the OPN4-expressing ganglion cells was the same among WT, MNU, and dKO mice (Figure 5).

The primary difference between the previous work and ours is the mouse models. Most of the previous work used mutant mouse models, including photoreceptor degenerating models (*rd/rd*), dysfunctional models (Cnga3-KO, Gnat1-KO), and retinal network mutant (mGluR6-KO). In contrast, we injected the MNU to eliminate photoreceptors in several days, limiting the risk of remodeling and compensatory effects. In our photoreceptors. Our novel observation of ipRGCs’ contribution to the PLR might be crucial for diagnosing eye disease in clinical situations. Furthermore, our observation will contribute to a more complete understanding of retinal circuitry from phototransduction to behavioral outcomes. Recently, ipRGCs’ functional significance has been re-evaluated (Paksarian et al., 2020; Chakraborty et al., 2022), and our work might contribute to finding new roles of ipRGCs in PLR.

### Limitation of the study

Jain et al. (2016) used MNU to examine the cellular contributions to the PLR. Although we conducted similar procedures, our results and conclusions do not agree with theirs. In their MNU-injected mice, pupil constriction in response to high light occurred only halfway compared to the WT mice, even though they observed an increased number of ipRGCs seven days after MNU injection. While MNU concentration, mouse strains, and light levels in their experiments were similar to our conditions, other conditions were distinct, including anesthesia (ours: no anesthesia, Jain et al: light anesthesia), and dark adaptation before PLR stimuli (ours: more than 30 min, Jain et al: 60-100 sec), which might cause the differential outcomes.

We conducted PLR measurements *in vivo* and ipRGC recordings *ex vivo* to investigate whether ipRGC light response explains the PLR in the MNU mice. We aligned the conditions in these measurements, such as light stimulus intensity and temperature. However, these conditions cannot be the same between *in vivo* and *ex vivo* experiments. Light intensity at the retina in the *in vivo* mouse eye might be dimmer due to light diffraction from the cornea and lens than in the MEA chamber during *ex vivo* studies. The mouse body temperature (38°C) is higher than the MEA recording chamber (33-34°C) and could explain the low percentage of fast responding ipRGCs during these studies. Even though some conditions differed, we found that both recordings exhibited similar results in MNU tissue, demonstrating that ipRGCs’ direct light response explains the robust PRL in the MNU mice.

## Acknowledgements

We would like to thank Dr. Dao-Qi Zhang for his advice on immunohistochemistry with melanopsin antibody, and Mr. Bashir Khatib-Shahidi for technical assistance.

## Author contributions

Conceptualization, T.I. and Z.S; Methodology, J.M.B and Z.J.S; Investigation, J.M.B., A.R.H., Z.J.S., M.K., M.A., Y.P., A.S., and T.I.; Writing – Original Draft, T.I.; Writing – Review & Editing, J.M.B., A.R.H., and T.I.; Funding Acquisition, J.M.B. and T.I.; Resources, T.I.; Supervision, T.I.

## Commercial relationships

none

